# Yanocomp: robust prediction of m^6^A modifications in individual nanopore direct RNA reads

**DOI:** 10.1101/2021.06.15.448494

**Authors:** Matthew T. Parker, Geoffrey J. Barton, Gordon G. Simpson

## Abstract

Yanocomp is a tool for predicting the positions and stoichiometries of RNA modifications in Nanopore direct RNA sequencing data. It uses general mixture models to identify differentially modified sites between two conditions, with good support for replicates. Yanocomp models across adjacent kmers and uses a uniform component to account for outliers, improving the accuracy of single molecule predictions. Consequently, Yanocomp can be used to measure modification stoichiometry, and correlate modifications with other RNA processing events.

**Availability:** Yanocomp is available under an MIT license at www.github.com/bartongroup/yanocomp.

## Introduction

Choices in RNA processing and modification determine mRNA coding potential and fate. Nanopore direct RNA sequencing (DRS) provides a unique insight into these processing outcomes using full-length reads representing single mRNA molecules (Garalde *et al*., 2018; Parker *et al*., 2020). Nanopore technology measures the changes in current caused by passing RNA molecules through a pore in a membrane. The measured signal depends on approximately 5 bases which block the narrow part of the pore at any one time (Garalde *et al*., 2018). Signals are translated into sequences using a neural network. This information can be used to examine features of mRNA processing including splicing and polyadenylation (Parker *et al*., 2020).

Nanopore DRS can also be used to map sites of RNA modifications such as N(6) methyladenosine (m^6^A), since modified bases cause characteristic changes in nanopore signal (Garalde *et al*., 2018). Several existing tools detect modifications using signal level data that has been aligned to a reference sequence using dynamic programming or hidden Markov models (HMMs) (Loman *et al*., 2015; Stoiber *et al*., 2016; Müller *et al*., 2019). These tools mostly rely on comparisons between wild type and samples with low levels of modification, e.g. from mutants lacking m^6^A (Parker *et al*., 2020; Leger *et al*., 2019; Pratanwanich *et al*., 2020). Differences between aligned signals can be identified using statistical methods such as a Kolmogorov-Smirnov test (Stoiber *et al*., 2016; Leger *et al*., 2019), or using a general mixture model (GMM), with two Gaussian components modelling modified and unmodified bases (Müller *et al*., 2019; Leger *et al*., 2019; Pratanwanich *et al*., 2020). The latter method has the advantage of also estimating the stoichiometry of modifications. Furthermore, fitted GMMs can be used to make single molecule predictions of modifications in individual reads. However, Gaussian components can be sensitive to outliers, which can skew parameters resulting in false positive results and lower quality single molecule predictions. This is an issue in nanopore DRS data since outliers can result from poor signal or alignment.

Here we present Yanocomp, a tool designed to identify RNA modifications from nanopore DRS signal data aligned using the HMM tool Nanopolish (Loman *et al*., 2015). Yanocomp models modified and unmodified bases using GMMs, and includes a uniform component which captures outliers, improving the robustness of the models. Yanocomp also predicts modifications in individual nanopore DRS reads, allowing correlation of modified bases with other RNA processing outcomes.

## Implementation

Yanocomp is implemented using scientific Python (Harris *et al*., 2020; Schreiber, 2017; Virtanen *et al*., 2020). It summarises the output of Nanopolish (Loman *et al*., 2015) into a table of mean current values per read, per position in each gene. Summarised tables are stored in an HDF5 file allowing random access of individual genes. Since 5 bases occupy the pore at any time, modified bases can cause deviations in current for at least 5 adjacent positions. We therefore use a 5-position sliding window to fit multivariate GMMs, comprising two Gaussian components with full covariance matrices. Modelling multiple positions improves the separation of modified and unmodified distributions. A third, uniformly distributed component improves the robustness of models to outliers, helping to prevent overfitting.

The fitting of GMMs is dependent upon parameter initialisation, which we perform using K-means with two clusters. Outliers are initially identified by calculating the median absolute deviation (MAD) of the Euclidean distances of each datapoint to the nearest cluster centroid, and selecting those above a distance of 0.5×MAD. GMMs are then fitted by expectation maximisation using pomegranate (Schreiber, 2017), with one model per biological replicate. Models from different replicates share the same underlying distributions but have independent weights for the contributions of each component. Gaussian distributions are labelled as modified or unmodified by comparison to existing models of unmodified kmers. Weights are used to estimate the number of modified and unmodified reads in each replicate. G-tests are then used to ensure homogeneity between replicates of the same condition and identify differences in modification rate between control and treatment conditions. Models can also be used to make single molecule predictions for each read.

## Example

We performed differential modification analysis of Arabidopsis mRNA using a mutant with low m^6^A levels (*vir-1*) and a complemented line (VIRc) (Parker *et al*., 2020). Yanocomp identified 20,033 modification sites, of which 16,389 (81.8%) had lower modification rates in *vir-1* (Figure 1A). An unbiased motif search at modification sites recovered the m^6^A consensus motif (Figure 1B). Modification sites were almost exclusively found in 3’UTRs, as expected in Arabidopsis (Figure 1C). Furthermore, comparison to miCLIP data (an m^6^A antibody cross-linking method) indicates that 84.1% of detected modification sites are within 5 nt of an orthogonally identified m^6^A site (Figure 1D).

**Figure 1:**
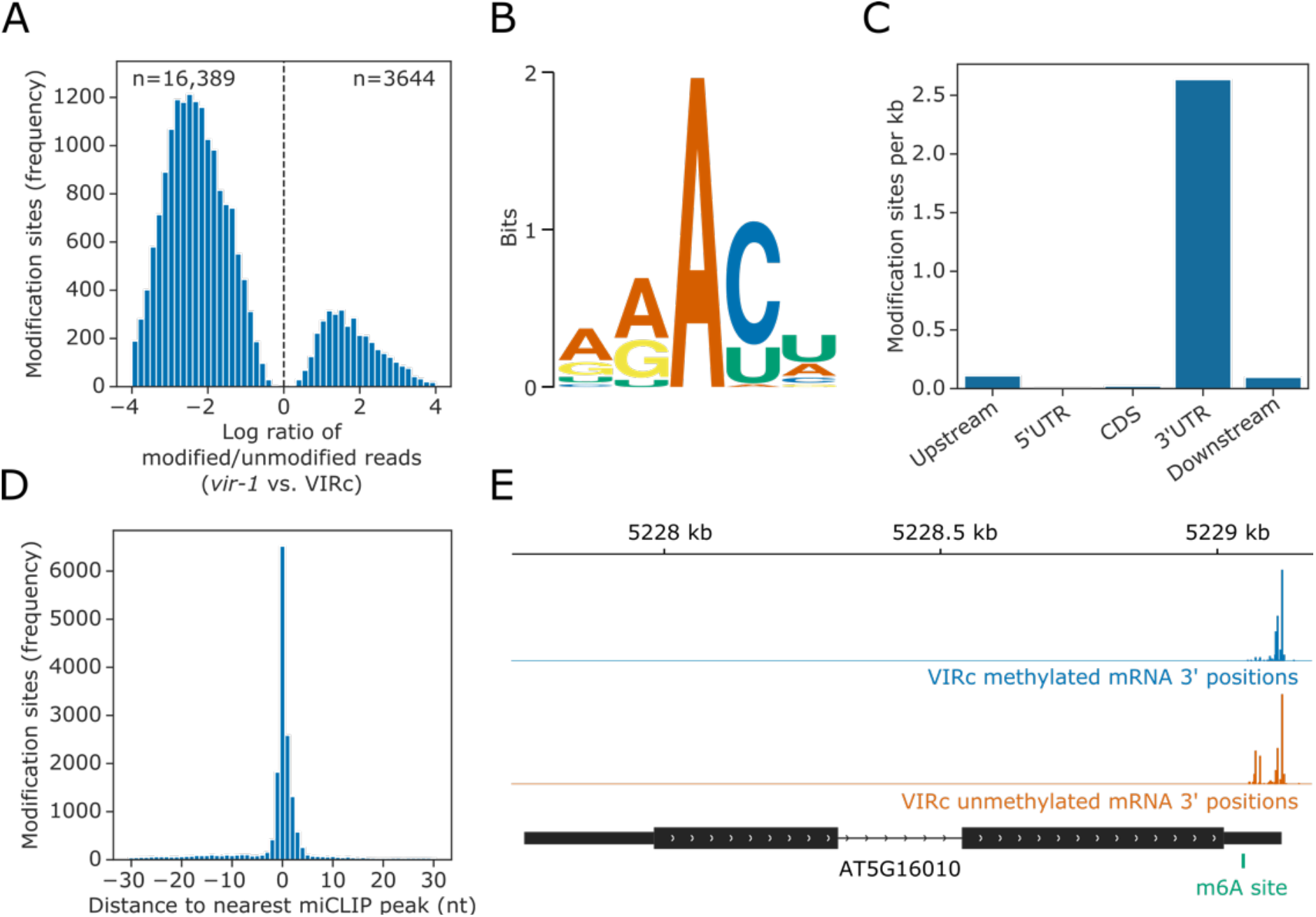
Yanocomp identifies m^6^A sites in *Arabidopsis thaliana*. **(A)** Histogram showing log ratio of modified/unmodified reads in *vir-1* compared to VIRc, for positions with significantly altered modification rates (FDR < 0.05). **(B)** Logo detected by MEME (Bailey *et al*., 2009) at positions with significantly altered modification rates. **(C)** Bar plot showing the number of modification sites per kilobase (kb) for different genic feature types of 48,149 protein coding transcript loci in the nuclear genome of the Araport11 reference. Upstream and downstream were defined as 200 nt regions before annotated transcription start sites and after annotated transcription termination sites, respectively. **(D)** Histogram showing the distribution of distances to the nearest miCLIP peak for each position with significantly altered modification rate. **(E)** Gene track showing 3’ positions for VIRc reads at the gene AT5G16010, which have been separated by modification status.

RNA methylation is linked to poly(A) site choice in Arabidopsis (Parker *et al*., 2020). To demonstrate the utility of Yanocomp single molecule predictions, we separated methylated and unmethylated reads in VIRc for the example gene AT5G16010 and compared 3’ end distributions. Unmethylated mRNAs had increased proximal polyadenylation compared to methylated mRNAs (p=7.9×10^−8^; Figure 1E). Therefore, Yanocomp predictions allow the phasing of RNA modifications with other processing outcomes.

## Conclusion

We have developed a tool that identifies differential RNA modifications using nanopore DRS signal. Yanocomp makes use of efficient data structures, fast modelling frameworks and multiprocessing to boost performance. We demonstrate that Yanocomp recovers genuine m^6^A sites in Arabidopsis, and that single molecule m^6^A predictions uncover interactions between RNA processing events. In future, we will use models of modified bases trained using Yanocomp to perform identification of m^6^A without requiring a low modification sample, for example by incorporation into HMMs.

